# Size matters – larger images are better remembered during free viewing

**DOI:** 10.1101/2020.05.24.113225

**Authors:** Shaimaa Masarwa, Olga Kreichman, Sharon Gilaie-Dotan

## Abstract

We are constantly exposed to multiple visual scenes, and while freely viewing them without an intentional effort to memorize or encode them, only some are remembered. It has been suggested that image memorability is influenced by multiple factors as depth of processing (1–3), familiarity (4), and visual category (5–7). However, this is typically investigated with intentional (8, 9) or unintentional (10, 11) encoding tasks. Furthermore, since visual memory relies on size-invariant visual perception (12–14), image size is not considered a contributing factor. Here we reasoned that during naturalistic free viewing of images without an encoding task, bigger images would be better remembered due to multiple factors (as vaster expanse of activated visual cortex (15, 16), deeper processing). In an extensive set of experiments participants (n=117) freely viewed small to large images (3°-24°) without any encoding task. Subsequently, they were presented with mid-sized images (50% already seen) and were asked to report if they recall seeing them or not. Larger images were better remembered (~23% more than smaller images), image memorability was proportional to image size, faces were better remembered, and outdoors the least. These were independent of image set, presentation order, or screen resolution. While multiple factors affect image memorability (1–7), here we show that during free viewing with no encoding task, a basic physical image dimension influences its memorability.

## Introduction

We are constantly exposed to many images, and despite not making intentional efforts, some of these images are burned into memory while others are not. It is yet unclear what determines image memorability under such unintentional (incidental) conditions. It is suggested that the “level of processing” of a stimulus modulates its memorability and this has been shown in the linguistic-lexical domain (e.g. when attending to fonts (“shallow”), words are less remembered than when attending to their semantic (“deeper”) aspects (2, 3)) and in the visual domain (e.g. when attending to gender faces are less remembered than when attending to honesty or likeableness (1)). Additional factors as familiarity and visual category also affect memorability (4, 5). Since high-level visual perception and its underlying mechanisms are assumed to be size invariant (12–14), image size is typically not considered as a factor influencing memorability. Here, in contrast to this view, we hypothesized that despite size invariance in visual perception, stimulus size may contribute to memorability such that larger images would be remembered better than smaller ones (Fig. 1A).

**Fig. 1.**
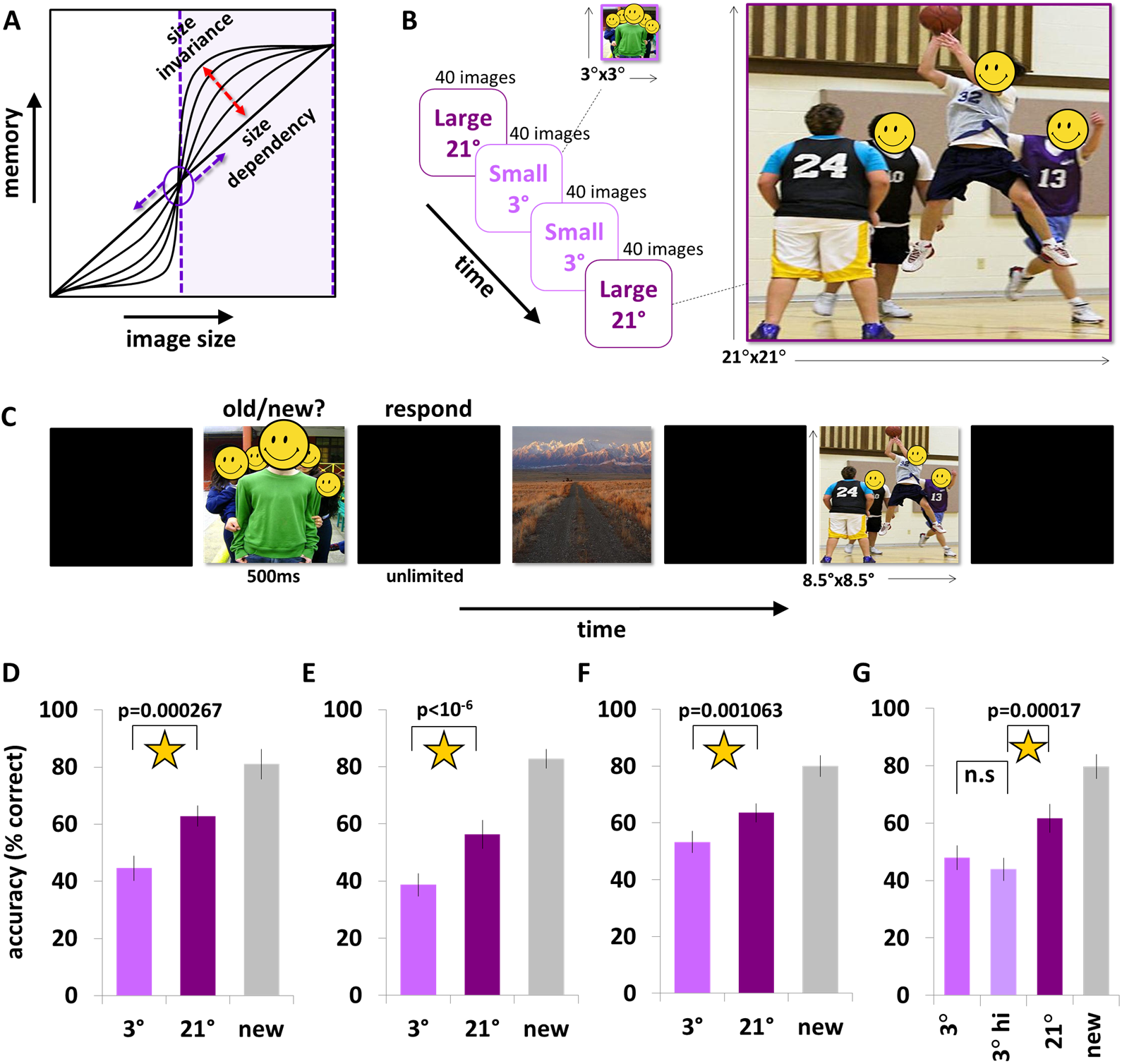
Larger images are remembered better than smaller ones during free viewing. (A) Hypothesized dependency of image memory on image size, ranging from size-dependency to size-invariance. (B) In Experiments 1-4 during the exposure phase participants viewed 80 small (3°x3°) and 80 large (21°x21°) images (2s each, small and large matched in contents) presented in 4 blocks of 40 images each. Participants were instructed to freely view the images without any memory or encoding instructions (see more details in text and Methods). (C) In Experiments 1-4 the test phase that followed the exposure phase included 320 midsized images (~8°x8°, 500ms each, 160 presented in the exposure), and participants were asked to report whether they recall seeing each image or not. (D)-(G) Test phase results reveal that in all experiments the images presented in larger format during the exposure phase were better remembered than the images presented in smaller format, and this was true in Experiment 1 (D), when we controlled for image sets in Experiment 2 (E), when we changed block order to control for recency or primacy effects(19–25) in Experiment 3 (F), or when we controlled for reduced resolution in Experiment 4 (G). Error bars represent SEM.

## Results

We tested this in an extensive set of five experiments (n=117). Each experiment started with passive viewing of images of different sizes (3° x 3° to 24° x 24°, exposure phase, Figure 1B) where participants were asked to freely view the images presented without being informed of any subsequent memory related task, and this was followed by an old/new recognition test phase with mid-sized images (mid-size determined by taking into account the cortical magnification factor (17, 18) (8° x 8°, Figure 1C)). In Experiment 1 participants (n=17) were exposed to 80 small (3° x 3°) and 80 large (21 ° x 21°) colored images that appeared in 4 blocks (large-small-small-large; each image presented for 2s, small and large images matched in contents, Figure 1B) and then were asked to make old/new memory judgements on a set of 320 mid-sized images (160 old, the old and new matched in contents, randomly ordered, 500ms exposure, 8° x 8° (midway between the small and large images when accounting for the cortical magnification factor (17, 18), see Supp. Mat.). Accuracy for larger (21°) images was significantly higher than for smaller (3°) images (Figure 1D, small: 44.6% ± 4.4% (SEM), large: 62.9% ± 3.7% (SEM), small vs. large p = 0.00027, t(15) = 4.73, paired 2-tailed). To rule out the possibility that the results may have been driven by specific images, a new group of participants (n=16) ran the same experiment but with the small and large image sets swapped (Experiment 2). Accuracy for larger images was again significantly higher than for smaller ones indicating that Exp. 1’s results were not driven by specific images (Figure 1E, small: 38.7% ± 4.1% (SEM), large: 56.4% ± 5.1% (SEM), p < 10^−5^, t(14) = 6.84, paired 2-tailed). To rule out the possibility that the results of Experiments 1 and 2 were driven by recency or primacy effects (19–25) as the larger images were presented in the first and last blocks, a new group of participants (n=17) underwent the same experiment but while swapping block order (small-large-large-small, Experiment 3). Here too, we found that accuracy was higher for the larger images regardless of block order (Figure 1F, small: 53.2% ± 3.9% (SEM), large: 63.5% ± 3.3% (SEM), p = 0.0011, t(15) = 4.04, paired 2-tailed). Since screen resolution may have degraded the information conveyed in the smaller images, in Experiment 4 (n=16), on top of the large (21°) and smaller images (3°) that were presented from the same viewing distance and spatial resolution as before (60cm), a new condition of the same viewing distance and spatial resolution as before (60cm) high resolution smaller images was included (3°). The high-resolution smaller images occupied the same visual angle as before (3° x 3°) but were based on large images presented from afar and thus while occupying the same visual angle provided much higher (x6.6^2^) spatial resolution. We found no difference between the accuracy in the small regular-resolution and the small higher-resolution conditions, and, as in Experiments 1-3, accuracy for larger images was higher than that of the smaller ones with higher resolution (Figure 1G, small (regular resolution, as in Exp. 1-3) 47.9% ± 4.2% (SEM), small (high-resolution) 43.9% ± 3.9% (SEM), large 61.7% ± 4.9% (SEM), small (regRes.) vs. small (highRes.) p > 0.168, t(14) = 1.45, small highRes. vs large p = 0.00017, t(14) = 5.058, paired 2-tailed), indicating that resolution was not likely to be the main factor influencing our results.

To parametrically investigate the effect of image size on memorability and the effect of visual categories on memorability we built a new 4 sizes (height and width of 3°, 6°, 12°, 24°) x 4 visual categories (faces, people, indoors and outdoors) experiment (Figure 2A). We used images with predefined memorability scores (LaMem Dataset (5)) such that image sets of each experimental size had equal memorability scores and equal contribution of each visual category (Figure 2A,B). Two new groups (n_1_=25, n_2_=26) underwent two versions of this experiment (image sets of size conditions were swapped across versions (3° with 24°, 6° with 12°, see Methods). Since there was no effect of experimental version and no interaction between size and version (Table S2) we collapsed the data across the 2 versions. We found a main effect of image size on memory (Figure 2C, F(3,150)=57.31 p<0.0001) with 3°<6°<12° (post-hoc p’s<0.0001, Table S2) indicating that in this range of image sizes (3°-12°) image memorability is size dependent (Fig. 2D). There was also a main effect of visual category on memory (F(3,150)=15.76 p<0.0001) with faces most memorable and outdoor scenes the least (Fig. 2E, Table S2). Image memorability scores (5) were significantly correlated with the actual memory performance (Fig. 2F, Table S3). Since each image appeared in one version as bigger and in the other as smaller, we examined for each image if size had any effect on its memorability by comparing performance between the two versions. This per-image analysis revealed that the images were significantly better remembered when they were presented in bigger format (Fig. 2G-I) and this was true for the big size difference (24° vs 3° mean accuracy difference of 23.4% ± 1.8% (SEM), t(79) = 13.0, p < 10^−20^, (n=80), see Fig. 2G,I) and for the smaller one (12° vs 6° mean accuracy difference of 9.04% ± 1.6% (SEM), t(79) = 5.58, p < 10^−6^, (n=80), see Fig. 2H-I). We also found that the highest accuracy across all experiments was to the new (unseen) images (see Table S1 and Figs. 1D-G, 2C,E), and no priming effects (implicit or explicit (26)) were found (Tables S4–S5).

**Fig. 2.**
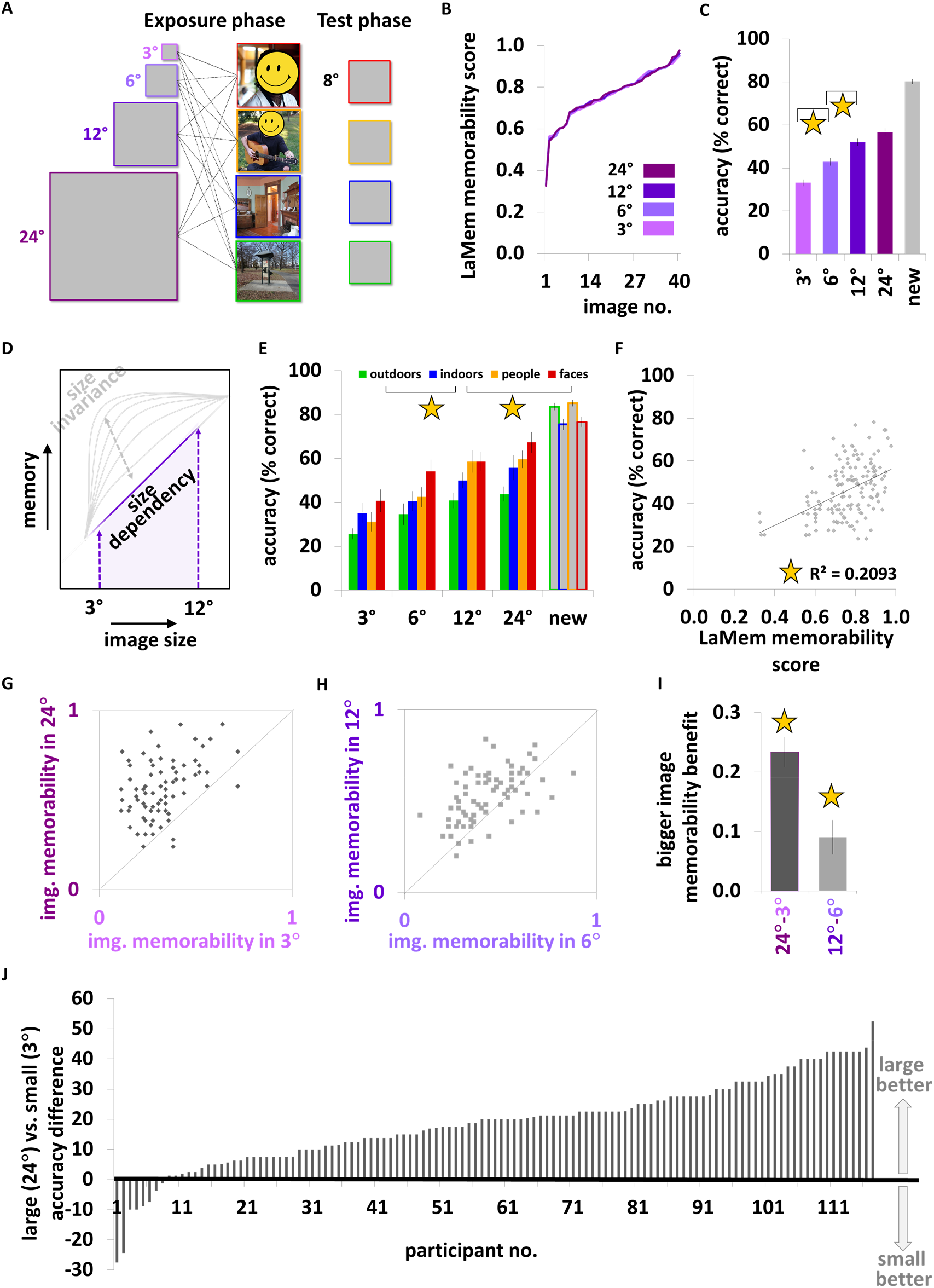
Image memorability during free viewing is linearly dependent on size for 3 ° – 12° images. (A) In Experiment 5 participants (n=51) viewed 4 blocks (random block order and within-block image order), each block included 40 uniformly sized images (either 3 °x3°, 6°x6°,12°x12° or 24°x24°, 2s/image, all from LaMem Dataset (5)) from 4 visual categories (faces, people, indoors and outdoors). Experimental instructions as in Experiments 1-4. In the test phase 320 mid-sized images (160 old, ~8°x8°, 500ms each) were presented. (B) Image memorability scores (5) were equally distributed across the experimental size conditions. (C) Significant effect of image size on image memory was found with images of 24° and 12° best remembered, 6° less, and 3° the least (Tables S4, S5). (D) Our results indicate that unintentional incidental memory (10, 11) of images is size-dependent for images of sizes 3°-12°.(E) Significant effect of visual category was found with faces and people best remembered, indoors less, and outdoors the least (Tables S4, S5). (F)-(I) Per-image analyses. (F) Image average accuracy across participants was significantly correlated with LaMem image memorability scores (Table S3). Images were better remembered if they were presented at 24° relative to 3° (G, I) or at 12° relative to 6° (H, I). (J) Large (21°/24°) vs. small (3°) image memory difference for all study participants (sorted by difference). Error bars represent SEM.

## Discussion

Our results show for the first time that physical dimension as image size has significant effect on unintentional (incidental (10, 11)) memory of images. This challenges the assumption that visual memory that relies on size-invariant visual perception is also size invariant. Note that we did not employ the classic incidental memory paradigm involving incidental encoding of images (e.g. 10, 11) or any intentional encoding (e.g. 8, 9). Rather, there was no encoding task and thus our results may not be comparable to earlier incidental or intentional memory studies (e.g. 9, 27–29). In addition, our old-new memory task on one item at a time may have further contributed to the consistently lower memory performance we found across our experiments relative to earlier encoding based findings (e.g. 8, 9, 30). In line with earlier studies (5–7), we found that face images were best remembered, and this was true across image sizes (limited to the categories and sizes we used), which could possibly be attributed to their social significance (31, 32). The size-memorability apparently linear relation we found in the size range that we investigated (3°-12°, Fig. 2C, D, and J for individual data (n=117)) may not generalize to bigger or smaller images. In fact we hypothesize that there would be a sharp drop in performance for images smaller than 3°, as well as for images that are “too big” to be perceived, as when viewing a movie from the first row in the cinema.

The size effect on memorability may be attributed to larger expanse of retinotopic cortex responding to bigger images (15, 16), possibly to different eye movement patterns, different spatial frequency contents across the different sizes, different spatial integration, attention, saliency or different levels of detail, all which may influence memorability. While our results may not generalize to active intentional or incidental memory paradigms, our study does demonstrate that physical image dimension can affect image memorability under conditions that closely mimic naturalistic daily visual behavior. However, it does not imply that other physical dimensions would have similar effects, or that other non-physical factors (e.g. cognitive) do not play a role in image memorability.

## Methods

### Participants

A group of 117 participants took part in the study, each participated in only one experiment (17 in Experiment 1 (12 women, aged 28.1 ± 6.9 year, 15 right handers), 16 in Experiment 2 (10 women, aged 25.4 ± 6.0 year, 15 right handers), 17 in Experiment 3(11 women, aged 25.1 ± 5.7 year, all right handers), 16 in Experiment 4 (10 women, aged 25.7 ± 6.3 year, 15 right handers), and 51 in Experiment 5 (26 in version 1, 25 in version 2, 31 women, aged 25.6 ± 6.6 year, 46 right handers). The experimental protocols were approved by the Bar Ilan University Ethics committee. All the participants signed a written informed consent before their participation. All participants had normal or corrected to normal far and near vision (all were checked for near and for far visual acuity before the experiment began).

### Experiments

All experiments were conducted in a dark room on an Eizo FG2421 24” HD LCD monitor with 1920×1080 pixels resolution running at 100 Hz (Experiment 4 was conducted on an Asus VG248QE 24” monitor with 1920×1080 pixel resolution running at 144 Hz) using an in-house developed platform for psychophysical and eye-tracking experiments (PSY) developed by Yoram S. Bonneh (33) running on a Windows PC. Viewing distance was 60cm from the screen in all experiments and conditions except for small highRes condition in Experiment 4 (see details in Experiment 4). Each experiment took approximately 25 min. ANOVA statistical analyses were performed with StatView software 5.0.

### Experiment 1

At the exposure phase, 160 images (coloured photographs) of the study set were presented in four blocks (each block of 40 images) in the following block order: Large Small Small Large (see Figure 1), image order within each block was randomized. Small images subtended a visual angle of 3.15°x3.15°, large images 20.78°x20.78°. Images were presented in a sequence, each image displayed for 2s followed by a 500ms black screen inter-stimulus interval, no response was required. Participants were asked to freely view and attend the images (fixation was not superimposed on the images, no fixation was required). All images were taken from the internet and resized to equal width and height (800×800 pixels) to avoid within- and across-condition size differences. These uniformly sized images were then scaled to be displayed according to the experimental condition (small or large). The images included different visual categories (faces, people, hands, animals, food, flowers, indoor places, outdoor places, and vehicles) and the images of each visual category were distributed equally between the small and large image sets.

The test phase followed the exposure phase, and participants were required to perform an old or new image memory task on 320 mid-sized images (visual angle of 8.39°x8.39°, see ‘Test phase image size’ below, 160 ‘old’ (previously seen in the exposure phase), 160 ‘new’) that were presented sequentially in random order. Each image appeared for 500ms followed by a black screen, participants were required to report if they recalled seeing each image (old) or not (new) without time limitations, no feedback was given.

### Experiment 2

To rule out the possibility that specific images may have biased the results of Experiment 1, in this experiment, the experimental protocol was precisely the same as Experiment 1 except that the image sets of the small and large conditions from Experiment 1 were switched (images displayed as small in Experiment 1 were now displayed as big and vice versa).

### Experiment 3

To rule out the possibility that the results of Experiments 1 and 2 may have been biased by recency or primacy effects (19–25) (as the large blocks appeared first and last), here we controlled for presentation order such that smaller image blocks appeared first and last. Images were assigned to blocks as in Experiment 1 but block order in the exposure phase was changed to Small Large Large Small. This allowed us to examine if size affects image memorability despite possible primacy or recency effects for small images.

### Experiment 4

In this experiment we wanted to rule out the possibility that the reduced memorability for smaller images may have resulted from degraded resolution for the smaller images relative to the larger images, especially given the screen pixel limitations. Therefore, in this experiment we included, as in Experiments 1-3, the small and large conditions from 60cm viewing distance, and in addition a new condition of small images with higher resolution (presented as large images but viewed from 4m to occupy the same visual angle as the original small images condition but provided much higher (x6.6^2^) spatial resolution). Specifically, the three conditions in the exposure phase included large images (visual angle of 20.78°x20.78°), small images (visual angle of 3.15°x3.15°), and small (higher resolution) images presented as big at 4m so that they occupy the same visual angle (same retinal image size) as the small images (3.15°x3.15°). We divided the original image set used in the exposure phase (as in Exp. 1-3, 159 out of the 160 images), to six subsets of similar size (3 with 26 images and 3 with 27 images), where each of these subsets had similar types of images (faces, people, places, food, etc.) and was used in one of the experimental blocks in the exposure phase. The exposure phase took approximately one minute longer since participants were required to move to a distant seating position for the small higher resolution condition. Participants responded via remote mouse across the experiment. Of the 16 participants that performed this experiment 6 participants underwent the Small Small(highRes) Large Small Small(highRes) Large block order, 6 participants underwent Small(highRes) Large Small Small(highRes) Large Small block order, and 4 performed the Large Small Small(highRes) Large Small Small(highRes) block order to counter balance condition order and so that each condition type would appear in a first block and in a last block of one of the versions. In the test phase 318 images were presented from 60cm at a visual angle of 8.39°x8.39° as in Experiments 1-3.

### Experiment 5

Here we wanted to parametrically investigate the effect of image size on memorability. In addition we also tested for visual categories’ effect on image memorability resulting in a four sizes (3°, 6°, 12°, 24° height/width) by four visual categories (faces, people, indoors and outdoors) design (Figure 2A). Images were taken from the “LaMem” database with per-image memorability scores (5) and were all resized to 900×900 pixels to avoid within- and across-condition size differences. These uniformly sized images were then scaled to be displayed according to the experimental condition (3°, 6°, 12°, 24°). We made sure that for each visual category (faces, people, indoors and outdoors) the memorability scores for that category across size conditions would be uniform (Figure 2B) and this assured that memorability scores across sizes would be comparable. A group of 25 participants ran version 1 of the experiment, 26 participants ran version 2. Image sets assigned to 3° condition in version 1 were assigned to 24° condition in version 2 and vice versa, image sets assigned to 6° condition in version 1 were assigned to 12° condition in version 2 and vice versa. Block order in each run (i.e. participant) regardless of version was randomized. The experiment was performed at a viewing distance of 60 cm.

The exposure phase included 160 images presented in 4 blocks, each of a fixed specific size (3°, 6°, 12°, or 24°) that included 40 images (10 from each visual category), block order and image order within blocks were random. As in Experiments 1-4, each image was presented for 2sec followed by a black screen of 500ms. Task was to view and attend the images. No response or fixation were required.

The test phase that followed included 320 mid-sized images (160 old, 160 new, visual angle of 8°x8°, see ‘Test phase image size’ below) that were presented sequentially in random order and participants were required to report for each image if they recall seeing it earlier (“old”) or not (“new”). Each image was presented for 500ms after which a black screen appeared until a response was given (there was no time limit). No feedback was given.

### Test phase image size

In order for the test phase images to be unbiased towards the small or large images, we aimed to present them at an intermediate size that would be midway (size wise) between the small and large images but in cortical rather than in retinal space. Taking into account the cortical magnification factor (CMF) that enhances foveal representations while reducing peripheral representations, we chose the midsize to be of a factor *a* bigger than the smaller images and a factor *a* smaller than the larger images such that 3° x *a*^2^ = 21° for Experiments 1-4 (resulting in test phase image size of 8°), and 3° x *a*^2^ = 24° for Experiment 5. Based on the estimates for M given in earlier studies (17, 18) we used 2 equations that estimate the magnification factor in human’s V1 (M_linear_ = 17.3/(E+0.75) by (17) and M_linear_ = 29.2/(E+3.67) by (18)), both yielding similar results for the test phase image size: 8.5° (Experiments 1-4) and 9.5° (for Experiment 5). Thus the choices of 8.39°x8.39° as the midsize for the test images in Experiments 1-4 and 8°x8° in Experiment 5 were slightly closer (in cortical space) to the size of the smaller images and thus unlikely to bias the results in favour of the bigger images.

## Acknowledgments

We would like to thank Yulia Golland, Nurit Gronau, Daniel Levy, Yoram Bonneh and Ifat Levy for discussions and suggestions, and Yuri Maximov for technical assistance. This work was supported by The Israel Science Foundation grant no. 1458/18 to SGD.

## Author contributions

SGD conceived the study, SGD and SM designed the study, SM prepared and ran the experiments, OK assisted in preparing the experiments, SM, SGD, and OK analyzed the data, SGD, OK and SM wrote the paper, all authors approved the submitted version of the paper.

## Supplementary Material

**Table S1.**
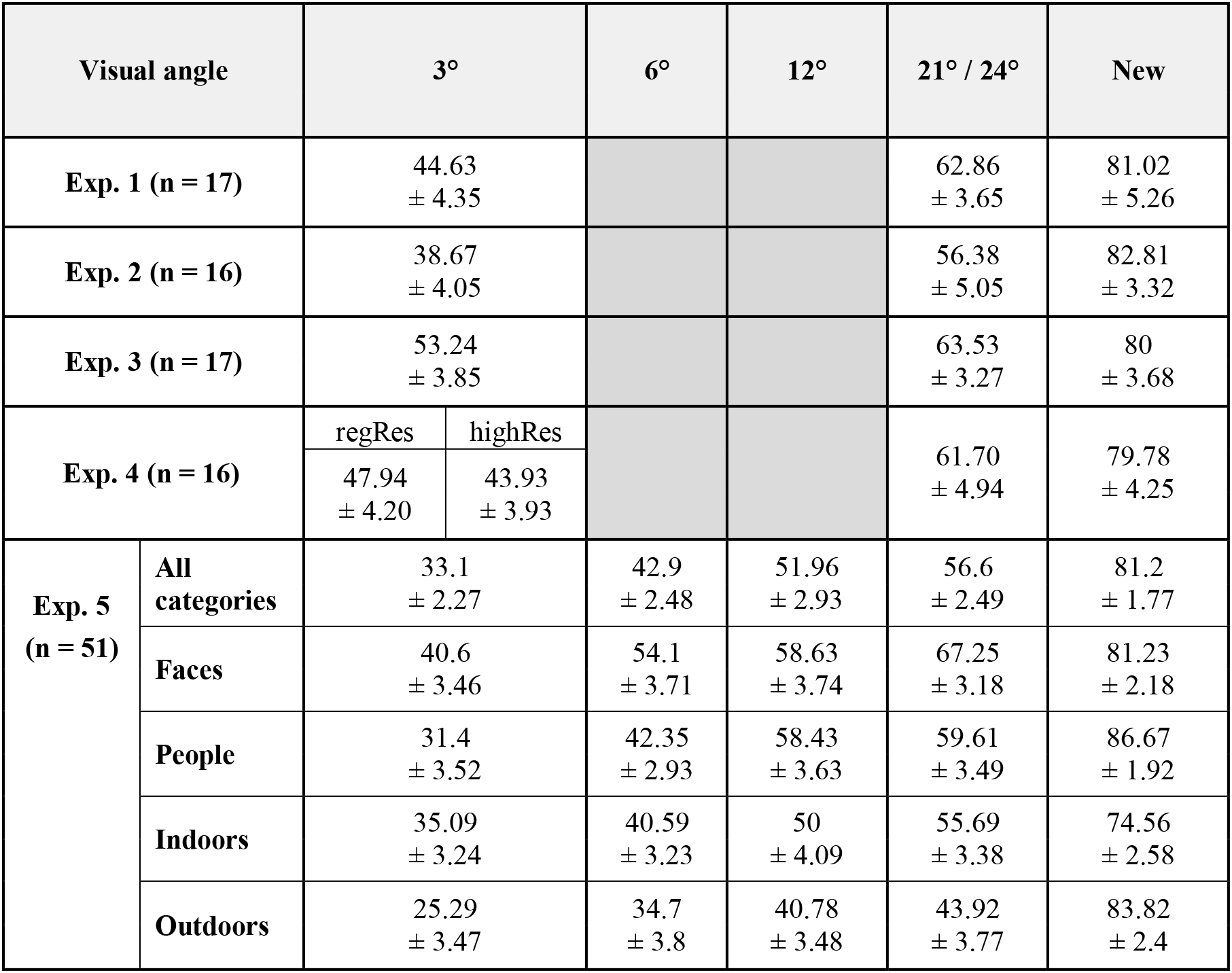
Accuracy performances – Experiments 1-5. For each experiment and each condition mean accuracy across participants ± SEM.

**Table S2.** Accuracy statistical analyses – Experiments 1-5. Accuracy analyses. Significant results appear in bold.

|  |  | T-test |  |  |  |
| --- | --- | --- | --- | --- | --- |
|  |  | Contrast | Results |  |  |
| Exp. 1 (n = 17) |  | 3°, 21° | t(15) = 4.732 **p = 0.0002** |  |  |
| Exp. 2 (n = 16) |  | 3°, 21° | t(14) = 6.85 **p = 10<sup>-6</sup>** |  |  |
| Exp. 3 (n = 17) |  | 3°, 21° | t(15) = 4.043 **p = 0.001** |  |  |
|  |  | ANOVA |  |  |  |
|  |  | Factors | Main effects | Interaction | Post-hoc<br>(Bonferroni/<br>Dunn) |
| Exp. 4 (n = 16) |  | Resolution<br>3°(regRes), 3°(highRes)<br>3°(highRes), 21° | F(3,45) = 21.13<br>**p < 0.0001** |  | p=0.425<br>**p=0.0009** |
| Exp. 5<br>(n = 51) | Version<br>X<br>Size | Version<br>(Version1, Version 2) | F(1,196) = 0.21<br>p = 0.646 | F(3,196)=0.12<br>p=0.94 |  |
|  |  | Size<br>(3°, 6°, 12°, 24°) | F(3,196) = 16.55<br>**p < 0.0001** |  |  |
|  | Size<br>X<br>Category | Size<br>3°, 6°<br>6°, 12°<br>12°, 24° | F(3,150) = 57.31<br>**p < 0.0001** | F(9,450)=1.84<br>p=0.0593 | p < 0.0001<br>**p < 0.0001**<br>p = 0.0177 |
|  |  | Category<br>Faces, People<br>Faces, Indoors<br>Indoor, Outdoors | F(3,150) = 15.76<br>**p < 0.0001** |  | p = 0.0101<br>**p = 0.0004**<br>**p = 0.0018** |

**Table S3.**
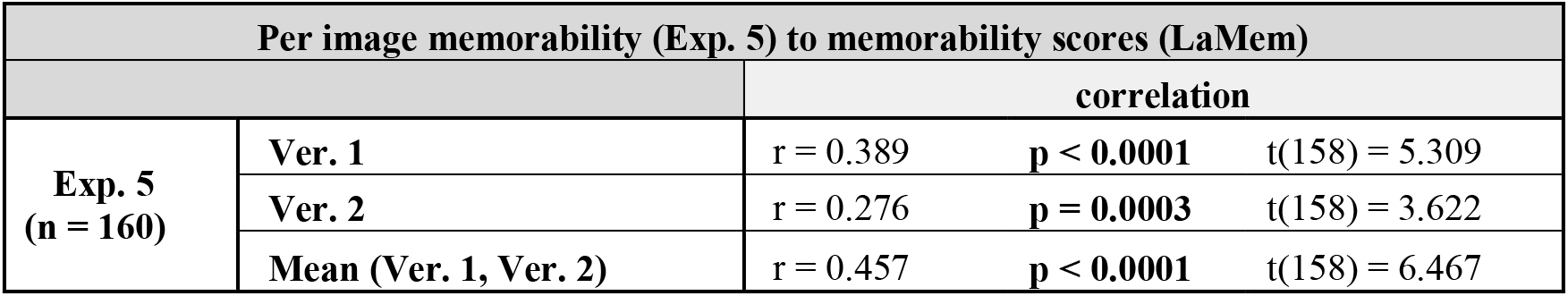
Experiment 5 per image analysis (n=160 images). For each image we calculated its average memorability in each version of Experiment 5 across all participants and compared these results to the LaMem image memorability scores (5). Since each image appeared in the 2 versions in 2 different sizes (either 3° and 24°, or 6° and 12°), for averaging across the versions (line 3), for each image we took the average memorability (mean across the 2 versions) as an approximation for its memorability for an average sized image (see Figure 2E).

**Table S4.** Reaction time performances – Experiments 1-5. For each experiment and each condition mean RTs across participants ± SEM. We did not find any explicit or implicit priming effects (see also Table S5).

| Visual angle | 3° |  | 6° | 12° | 21° / 24° | New |
| --- | --- | --- | --- | --- | --- | --- |
| Exp. 1 (n = 17) | 1041.62<br>± 53.48 |  |  |  | 984.5<br>± 41.96 | 947.647<br>± 33.33 |
| Exp. 2 (n = 16) | 1056.53<br>± 53.80 |  |  |  | 1005.72<br>± 50.96 | 1005<br>± 62.99 |
| Exp. 3 (n = 17) | 954.56<br>± 41.58 |  |  |  | 934.412<br>± 40.69 | 943.12<br>± 38.39 |
| Exp. 4 (n = 16) | regRes | highRes |  |  | 1071.78<br>± 61.36 | 1112.84<br>± 66.97 |
|  | 1101.21<br>± 73.99 | 1118.56<br>± 66.48 |  |  |  |  |
| Exp. 5 (n = 51) | 1055.16<br>± 261.8 |  | 1041.8<br>± 241.2 | 1003.8<br>± 218.0 | 1034.1<br>± 256.8 | 1044.8<br>± 237.7 |

**Table S5.** Reaction time statistical analyses – Experiments 1-5. Significant effects are indicated in bold. Note that in the 2 significant findings, the larger images showed faster RTs. Further analyses to examine possible priming effects (26) ((i) whether RTs of old images were faster than RTs of new images, and (ii) whether RTs of correctly remembered old images were faster than RTs of non-remembered old images) did not yield any significant findings with the mean or with the median RT in any of the experiments.

|  |  | T-test |  |  |
| --- | --- | --- | --- | --- |
|  |  | Contrasts | Results |  |
| Exp. 1 (n = 17) |  | 3°, 21° | t(15) = 2.380 | **p = 0.031** |
| Exp. 2 (n = 16) |  | 3°, 21° | t(14) = 2.186 | **p = 0.046** |
| Exp. 3 (n = 17) |  | 3°, 21° | t(15) = 1.025 | p = 0.321 |
|  |  | ANOVA |  |  |
|  |  | Factors | Main effects | Interaction |
| Exp. 4 (n = 16) |  | Resolution | F(3,45) = 1.321<br>p = 0.279 |  |
| Exp. 5<br>(n = 51) | Size<br>X<br>Category | Size | F(3,150) = 1.859<br>p = 0.139 | F(9,450) = 0.67<br>p = 0.7365 |
|  |  | Category | F(3,150) = 0.212<br>p = 0.8883 |  |

## References

1. G. H. Bower, M. B. Karlin, Depth of processing pictures of faces and recognition memory. J. Exp. Psychol. 103, 751–757 (1974).

2. F. I. M. Craik, E. Tulving, Depth of processing and the retention of words in episodic memory. J. Exp. Psychol. Gen. 104, 268–294 (1975).

3. M. Moscovitch, F. I. M. Craik, Depth of processing, retrieval cues, and uniqueness of encoding as factors in recall. J. Verbal Learning Verbal Behav. 15, 447–458 (1976).

4. A. D. Wagner, J. D. E. Gabrieli, M. Verfaellie, Dissociations between familiarity processes in explicit recognition and implicit perceptual memory. J. Exp. Psychol. Learn. Mem. Cogn. 23, 305–323 (1997).

5. A. Khosla, A. S. Raju, A. Torralba, A. Oliva, Understanding and Predicting Image Memorability at a Large Scale in 2015 IEEE International Conference on Computer Vision (ICCV), (IEEE, 2015), pp. 2390–2398.

6. W. Sato, S. Yoshikawa, Recognition memory for faces and scenes. J. Gen. Psychol. 140, 1–15 (2013).

7. P. Isola, J. Xiao, D. Parikh, A. Torralba, A. Oliva, What makes a photograph memorable? IEEE Trans. Pattern Anal. Mach. Intell. 36, 1469–1482 (2014).

8. T. F. Brady, T. Konkle, G. A. Alvarez, A. Oliva, Visual long-term memory has a massive storage capacity for object details. Proc. Natl. Acad. Sci. 105, 14325–14329 (2008).

9. L. Standing, J. Conezio, R. N. Haber, Perception and memory for pictures: Single-trial learning of 2500 visual stimuli. Psychon. Sci. 19, 73–74 (1970).

10. C. C. Williams, J. M. Henderson, F. Zacks, Incidental visual memory for targets and distractors in visual search. Percept. Psychophys. 67, 816–827 (2005).

11. I. S. Utochkin, J. M. Wolfe, Visual search for changes in scenes creates long-term, incidental memory traces. Attention, Perception, Psychophys. 80, 829–843 (2018).

12. K. Grill-Spector, et al., Differential Processing of Objects under Various Viewing Conditions in the Human Lateral Occipital Complex. Neuron 24, 187–203 (1999).

13. E. T. Rolls, G. C. Baylis, Size and contrast have only small effects on the responses to faces of neurons in the cortex of the superior temporal sulcus of the monkey. Exp. Brain Res. 65, 38–48 (1986).

14. Y. Han, G. Roig, G. Geiger, T. Poggio, Scale and translation-invariance for novel objects in human vision. Sci. Rep. 10, 1–13 (2020).

15. I. Levy, U. Hasson, G. Avidan, T. Hendler, R. Malach, Center–periphery organization of human object areas. Nat. Neurosci. 4, 533–539 (2001).

16. R. B. H. Tootell, et al., The Retinotopy of Visual Spatial Attention. Neuron 21, 1409–1422 (1998).

17. J. C. Horton, W. F. Hoyt, The Representation of the Visual Field in Human Striate Cortex. Arch. Ophthalmol. 109, 816 (1991).

18. P. Kovács, B. Knakker, P. Hermann, G. Kovács, Z. Vidnyánszky, Face inversion reveals holistic processing of peripheral faces. Cortex 97, 81–95 (2017).

19. R. A. Bjork, W. B. Whitten, Recency-sensitive retrieval processes in long-term free recall. Cogn. Psychol. 6, 173–189 (1974).

20. M. W. Howard, V. Venkatadass, K. A. Norman, M. J. Kahana, Associative processes in immediate recency. Mem. Cognit. 35, 1700–1711 (2007).

21. D. Rundus, Analysis of rehearsal processes in free recall. J. Exp. Psychol. 89, 63–77 (1971).

22. H. Shteingart, T. Neiman, Y. Loewenstein, The role of first impression in operant learning. J. Exp. Psychol. Gen. 142, 476–488 (2013).

23. W. A. McKenzie, M. S. Humphreys, Recency effects in direct and indirect memory tasks. Mem. Cognit. 19, 321–331 (1991).

24. E. Capitani, S. Della Sala, R. H. Logie, H. Spinnler, Recency, primacy, and memory: reappraising and standardising the serial position curve. Cortex 28, 315–342 (1992).

25. I. Fischler, D. Rundus, R. C. Atkinson, Effects of overt rehearsal procedures on free recall. Psychon. Sci. 19, 249–250 (1970).

26. E. Tulving, D. Schacter, Priming and human memory systems. Science (80-.). 247, 301–306 (1990).

27. C. C. Williams, Incidental and intentional visual memory: What memories are and are not affected by encoding tasks? Vis. cogn. 18, 1348–1367 (2010).

28. T. F. Brady, T. Konkle, G. A. Alvarez, A review of visual memory capacity: Beyond individual items and toward structured representations. J. Vis. 11, 4–4 (2011).

29. J. M. Wolfe, Y. I. Kuzmova, How many pixels make a memory? Picture memory for small pictures. Psychon. Bull. Rev. 18, 469–475 (2011).

30. R. N. Shepard, Recognition memory for words, sentences, and pictures. J. Verbal Learning Verbal Behav. 6, 156–163 (1967).

31. A. Todorov, C. P. Said, A. D. Engell, N. N. Oosterhof, Understanding evaluation of faces on social dimensions. Trends Cogn. Sci. 12, 455–460 (2008).

32. A. Todorov, N. Oosterhof, Modeling social perception of faces. IEEE Signal Process. Mag. 28, 117–122 (2011).

33. Y. S. Bonneh, Y. Adini, U. Polat, Contrast sensitivity revealed by microsaccades. J. Vis. 15, 11 (2015).

